# Mandrake: visualising microbial population structure by embedding millions of genomes into a low-dimensional representation

**DOI:** 10.1101/2021.10.28.466232

**Authors:** John A. Lees, Gerry Tonkin-Hill, Zhirong Yang, Jukka Corander

**Affiliations:** MRC Centre for Global Infectious Disease Analysis, School of Public Health, Imperial College London, London, United Kingdom; Department of Biostatistics, University of Oslo, Norway; Norwegian University of Science and Technology; Aalto University; Parasites and Microbes, Wellcome Sanger Institute, Cambridge, UK; Helsinki Institute for Information Technology HIIT, Department of Mathematics and Statistics, University of Helsinki, Finland

**Keywords:** pathogens, population structure, visualisation, genomics, dimensional reduction

## Abstract

In less than a decade, population genomics of microbes has progressed from the effort of sequencing dozens of strains to thousands, or even tens of thousands of strains in a single study. There are now hundreds of thousands of genomes available even for a single bacterial species and the number of genomes is expected to continue to increase at an accelerated pace given the advances in sequencing technology and widespread genomic surveillance initiatives. This explosion of data calls for innovative methods to enable rapid exploration of the structure of a population based on different data modalities, such as multiple sequence alignments, assemblies and estimates of gene content across different genomes. Here we present Mandrake, an efficient implementation of a dimensional reduction method tailored for the needs of large-scale population genomics. Mandrake is capable of visualising population structure from millions of whole genomes and we illustrate its usefulness with several data sets representing major pathogens. Our method is freely available both as an analysis pipeline (https://github.com/johnlees/mandrake) and as a browser-based interactive application (https://gtonkinhill.github.io/mandrake-web/).

## Introduction

Advances in DNA sequencing technology have recently made whole-genome sequencing both affordable and scalable enough for routine use in pathogen surveillance by research organizations and public health agencies around the world (1,2). A striking example of this is genomic surveillance of the SARS-CoV-2 virus for which over one million genome sequences became available in just 15 months after its initial discovery (3). To shed light on population genomic data at this scale calls for new tools that can be used for rapid exploration of the structure among the samples, with particular emphasis on detecting clusters of similar sequences (4,5). Many species do not have good quality schemes to label input genomes, or suffer from poor quality or missing metadata, so unsupervised methods are of particular interest when exploring data (6).

An additional challenge to the large number of individual genomes arises from the fact that genomic datasets typically have a very large number of features, for example when using SNPs or k-mers to represent sequence variation, each sample may typically have 10^6^ -10^8^ such markers. These markers are frequently used to calculate genetic distances between samples, the number of which grows as the number of samples squared, such that one million samples will have of the order of 10^11^ distances between them. Such high dimensionality of population genomic data is beyond the capability of most analysis methods available today, rendering it difficult to gain insight into the data structure in a fast and robust manner. In this paper we explore and extend a class of methods which aims to reduce the dimensionality of such data to only two dimensions, in a manner which supports ready visualization and identification of clusters.

An embedding seeks to find a lower-dimensional representation of data where the distances in the lower dimensional space *y* (output) are an accurate representation of distances in the higher dimensional space *x* (input). Intuitively, genetically similar samples should be close together in the embedding space, and genetically distant samples should be further apart in the embedding space. Embedding spaces may be linear combinations of the input dimensions as in principal component analysis and multidimensional scaling, but here we focus on non-linear methods, which can infer potentially complex manifolds relating input to output spaces in an unsupervised data-driven manner. This means, unlike in linear methods, the transform in one part of the input space may be quite different to another part of the space.

One such method is t-distributed stochastic neighbour embedding (t-SNE) (7,8). Rather than minimising a distance between the input and output data, t-SNE minimises the Kullback-Leibler divergence between two probability distributions defined by the input and output data. The input conditional probability distribution p_j|i_ between a pair of samples *i* and *j* is given by:

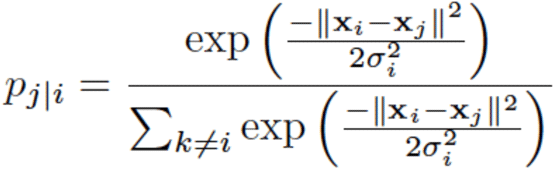

which equals the probability that x_i_ would pick x_j_ as its neighbour when sampling from a normal probability distribution centred at x_i_ with variance *σ*_*i*_. To define this probability, it is necessary to set a ‘perplexity’ which can be interpreted as the expected number of neighbours for each sample. Lower values of perplexity favour more local structure, whereas higher values assign greater weight to the global structure. Given the desired perplexity level, the variances and the corresponding conditional probabilities can be computed for each sample efficiently and in parallel (9), a technique known as entropic affinity.

In the output space t-SNE defines the probabilities q_ij_ using a student t-distribution with one degree of freedom (a Cauchy distribution):

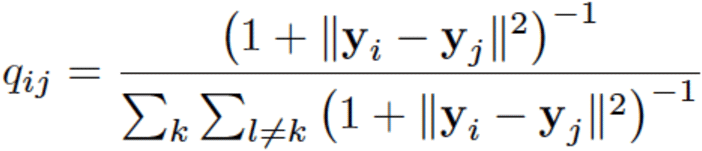

The use of a heavy tailed distribution rather than a normal distribution allows points to be further apart without affecting the divergence too much, and is also faster to compute.

A popular measure of discrepancy between two probability distributions P(x) and Q(x) is given by the Kullback–Leibler divergence, which is defined as:

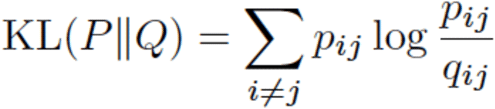

The t-SNE algorithm minimises this divergence iteratively, thus giving an embedding *y* with a probability distribution for between sample distances which is as similar as possible to the probability distribution for between sample distances in the higher dimensional data *x*.

t-SNE and related methods have been used extensively to represent and visualise data from numerous fields of research and they have recently been considered for analyzing population structure in both human and pathogen populations, as well as data from single-cell genomics (10–13). As these are unsupervised methods, they do not use sample labels to find the embedding. Due to the choice of the output probability distribution, distances between local samples are preserved, whereas global distances are less well preserved. Consequently, t-SNE is often used to identify clusters in high-dimensional data, which may correspond to units of population structure such as species, strains or lineages. Alternatively they may map onto sample labels, such as their geographical origin or cell type.

However, t-SNE is not optimising the embedding to find clusters. So, when clusters do emerge, they are an indirect consequence of preserving local structure in the data. The recently developed method of stochastic cluster embedding (SCE) (14) generalises t-SNE to include an additional scaling parameter, replacing the denominator of q_ij_ in the Kullback-Leibler divergence. The authors of this method show that this scale factor can be chosen to exactly replicate t-SNE, or alternatively can be tuned to effectively increase the ‘repulsion’ between points, targeting distinct clusters forming in the output embedding, which are easier to visualise and interpret.

In this paper we extend the SCE method to use a variety of genomic data modalities as input, improve its performance on large datasets, and add a range of output visualisations. Our method allows users to rapidly gain insights into structure present in very large genome datasets, which we show corresponds well with model-based genetic clustering algorithms. We implemented our method as a piece of open-source software called mandrake (https://github.com/johnlees/mandrake), and as a static web application (https://gtonkinhill.github.io/mandrake-web).

## Methods

### Calculating between sample distances from genome data

As input, mandrake takes one of three types of data: a multiple sequence alignment, a set of k-mer sketches (can be created from assembled or sequence read data), or a binary presence absence matrix (which is typically used to represent genes, but can be used to represent other genetic elements). These are all treated in fundamentally the same way, as feature matrices, with *N* samples along rows, and *M* features (SNPs, k-mers or genes) along columns. Although typically genomic datasets have been ‘wide’, with many more features than samples, the scale of data means this is no longer the case, and we are now able to analyse the case with more samples than genomic features.

To calculate input distances **X** from the feature matrix **A** we can compute **X** = *M* - **AA**^T^, which counts the number of shared features between every pair of samples (the similarity) and converts this to a distance by subtracting from the maximum shared features *M*. This is a symmetric matrix with zeros on the diagonal. We note that more sophisticated genetic distance calculations are possible by accounting for base frequencies and varying transition rates between classes, but we do not consider such distances here.

A difficulty is that both the number of calculations needed to find **X** and the amount of memory to store **X** grows as *N*^2^. Here we use methods which are fast enough to scale to *N*^2^ for at least one million samples, but such a matrix would still require at least 2Tb of memory (or disk space). To avoid this major resource issue, we cut the size of **X** down using one of two methods. The first is to set a distance threshold above which entries from **X** are discarded. The second, which we use for all analyses here, is to retain just the *k* nearest neighbours for each sample (excluding self distances, and including any ties). This means **X** grows linearly in size with *Nk* in a predictable way, making memory allocations efficient. As the perplexity sets the expected number of neighbours, choosing a *k* above the desired perplexity will typically give good results. In practise, we store **X** as a sparse matrix in coordinate (‘triplet’) format, with three ordered lists of *i, j* and *x*_*ij*_ for each retained distance. We save these to disk so they can be reused by other programs, or by mandrake to re-run the embedding without recomputing distances.

When **A** is a multiple sequence alignment, we code each row using the four DNA bases, each in its own dynamic bitset with the same length as the alignment, storing 1 if the base is present in that sample at that position, and 0 otherwise. Elements *x*_*ij*_ *= M* – *∑a*_*im*_*a*_*jm*_ are then computed by ANDing each of the four bitsets and counting the total number of bits that are on (popcount). The use of bitsets ensures efficient packing into 64-bit words, which makes the boolean AND operation and subsequent popcount very fast to complete across all *M* sites. If **A** is a gene presence/absence matrix, the procedure is similar, but only a single bitset is needed for each gene.

For sequence assemblies or sequence reads, which are unaligned, we count the number of shared k-mers between samples. Reads can be ‘cleaned’ by first removing low frequency k-mers, which typically are a consequence of sequencing error. Rather than using all k-mers, of which there are a prohibitively large number (15), we use a ‘sketching’ approach pioneered by the popular mash software, which instead uses a hash function [a hash function here transforms a k-mer sequence to a 64-bit integer] to uniformly subsample a fixed-size subset of the total k-mers (16). The proportion of shared k-mers (the Jaccard distance) can be computed by the size of the intersection of the retained hashes. We use two further modifications to this process. First, we use the method of bindash (17) to bin hashes and calculate distances between them (which turns out to be very similar to the dynamic bitset approach, but using bits of the calculated hash instead of DNA bases). Secondly, we optionally enable the approach of PopPUNK which calculates the Jaccard distance at multiple k-mer lengths and regresses their depletion at longer lengths to calculate core and accessory distances within a species (18). In practise we use PopPUNK’s sketching and distance library pp-sketchlib (https://github.com/johnlees/pp-sketchlib) which optimises sketching and distance calculation from assembly or read data, and has an API which can be directly called from python.

The computation of each row of **A** and reduction to the *k* nearest neighbours is embarrassingly parallel across up to *N* processes. We use OpenMP to achieve CPU parallelism. pp-sketchlib can also make use of CUDA compatible GPUs for further parallelism.

To convert distances in **X** to conditional probabilities **P** we used the entropic affinity using interval bisection to find a suitable variance given a user-input perplexity parameter (9). We used the implementation in scikit-learn, adding CPU parallelism with OpenMP (19).

### Stochastic cluster embedding (SCE)

We give a brief overview of the mechanism behind SCE, but note that full details are covered in the original publication (14). We also based our implementation on the reference implementation available at https://github.com/rozyangno/sce, and note the main changes here.

The main difference between the SCE algorithm and the t-SNE algorithm described above stems from the scaling factor *s* which appears in the denominator of *q*_*ij*_. In SCE, an alternative choice of *s* is determined which makes clusters apparent in visualisations (determined by a user study in the original publication). This allows the objective function to be minimised, *D* (the modified Kullback-Leibler divergence):

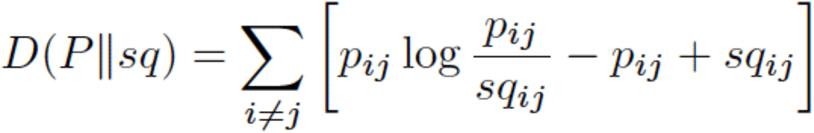

to be written in terms of an attraction, repulsion and constant with respect to *q* | *s*:

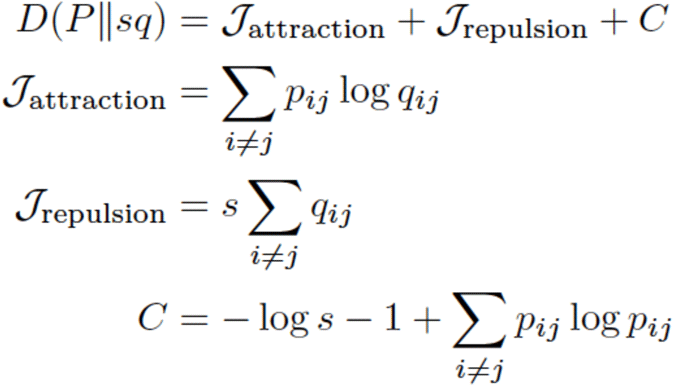

The stochastic cluster embedding method optimises *D* using stochastic gradient descent (SGD), a popular method to fit neural networks (20). Here, the output embedding **Y** is updated given the current s, then s is recomputed using the update **Y**. This is repeated for a specified number of iterations, chosen such that *D* reaches a stable minima. To stochastically update **Y**, a pair of samples *i, j* are chosen at random in proportion to their conditional probabilities *p*_j|i_, and the gradient ▽ of their attraction term calculated (such that *C* can be ignored). Then, a pair of samples i, j are chosen at random and the gradient of their repulsion term calculated. In SGD a learning rate ηis used to update **Y** by making a small step down the direction of the gradient *y*_*i*_ ← *y*_*i*_ – η_*t*_ ⍰ at iteration *t*. The learning rate decreases across the total *T* iterations *T* as 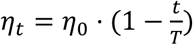. Larger steps are taken in early iterations, and smaller steps are taken in later iterations closer to convergence. **Y** is initialised by drawing *y*_*i*_ ∼ *U*(0,10−4) along each dimension.

While an additional drawback of t-SNE was that the iterative optimisation is challenging to directly scale to larger datasets, SGD is simpler to parallelise. At each step updating **Y**, *w* workers can independently pick pairs of points *i, j* to update. Ideally for CPU parallelism, *w* will be chosen equal to the number of physical cores, and for GPU parallelism w will be chosen to be large (10^5^ or more) to maximise device occupancy. A potential issue arises if two workers try to update the same *i* or *j* at the same time (bearing in mind the additional complication that these workers may not be in sync). This becomes more likely when the number of active workers is not much less than the number of samples. We address this in the CPU implementation by using atomic operations to preserve memory integrity, and when overwritten by another worker, retry with another pair. CUDA global memory is not directly affected by memory integrity issues from race conditions, but we still use an atomic operation to update **Y** rather than a simple overwrite. In each case, as long as memory integrity is preserved, the stochastic nature of the algorithm will correct for missteps in subsequent iterations, as long as they do not dominate. Additionally, while atomic operations are faster than locks, they become slower when multiple threads are attempting to operate on the same memory address, leading to a reduction in efficiency. We therefore output the proportion of workers found to be ‘clashing’ at each iteration, so users are aware they may wish to lower *w* when analysing smaller *N*.

We also note that we use the method of Walker (21) for drawing discrete random variables to precompute tables to draw edges from *P*_*j*|*i*_ in constant time, reimplementing the GSL library implementation in C++ (22). We also use the fast parallel random number generator from the dust package (23), which is based on the xoshiro128+ generator (24), and can be used to produce uncorrelated pseudorandom 32-bit integers in parallel on both CPUs and GPUs. This also removed all link time dependencies from the compiled code, which made compilation into WebAssembly straightforward (see below).

### Visualising embeddings

We automatically output the final embedding **Y** in four formats:

- A simple text file with N rows and two columns, for reuse by other programs or plotting software. A separate file listing sample names, and optionally clusters, is also created.
- An interactive HTML plot using the WebGL mode of plot.ly (25). This can be viewed in a web browser, and scales up to millions of points. Embedding positions and labels appear on hover. For smaller datasets sample names also appear on hover, but this can be turned off (as resulting files can be extremely large on disk).
- A static image using matplotlib (26).
- A .dot network file, which can be loaded for interactive viewing along with sample labels in Microreact (27).

To add colour to samples in the plot, the user can either provide labels, or labels can be generated by performing a spatial clustering on the embedding. For the latter, we use HDBSCAN, as this usually works well on well-separated clusters of unspecified shape. We centre and normalise the embedding to [-1, 1] in each direction, use a minimum cluster size of two, and minimum distance between clusters of 0.02 (28). HDBSCAN may label some points as ‘noise’, which are useful for potential singleton clusters, though care should be taken not to group noise points into a cluster.

Colours for classes are chosen by randomly sampling from RGB space. We tried selecting from HSL or HSLuv space, which are perceptually uniform colour spaces to the human eye, but found empirically that contrast between labels was poorer than from RGB colours. We found that for many of the genomic datasets we ran mandrake on, well-separated clusters were a common feature (for example separating species). In the embedding output this leads to many points overlapping, and although clusters can clearly be identified, their size is obscured. To help remedy this, we included an additional (static) hexagon density plot which shows a heatmap of the number of samples in each region of the plot.

We also include code to create a video of the embedding process as the SGD algorithm runs, which is particularly useful for monitoring convergence. We take the current embedding and objective function at 400 points across the total number of iterations, create a static plot, and use these as frames in the output animation (at 20fps, so videos are 20s in duration). In the CUDA code, the copy of the current embedding is launched asynchronously to the main SGD kernel run, so it has a negligible impact on run time. We optionally add sound by mixing decaying triangular wave oscillators at a frequency proportional to the maximum movement along each dimension between each frame. This sound is in stereo, with each channel corresponding to an SCE dimension.

Initially our code sampled frames uniformly from the SGD iterations, however this led to animations where at the start points moved too fast, and at the end too slow. This is due to the decreasing learning rate η. We decided instead to sample uniformly from the total amount of learning completed, so when more learning (and larger changes to the embedding) was being done more frames would be taken, and when less learning (and smaller changes to the embedding) was being done fewer frames would be taken.

As we use a linearly decreasing learning rate, learning grows quadratically, so we sample proportional to its inverse (the square root). More formally, the total amount of learning at iteration κ ≤*T*is given by:

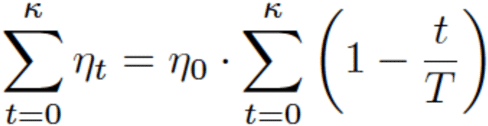

which can be approximated by an integral (ignoring a small constant term as T >> 1):

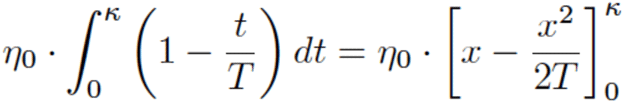

The total amount of learning completed when κ *=T* is therefore 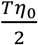 which can be subdivided equally into *f* frames, which can be done by taking a sample at iteration κif the total learning is an integer multiple *c* of 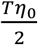:

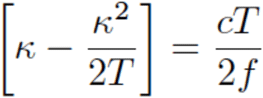

rearranging to find κ:

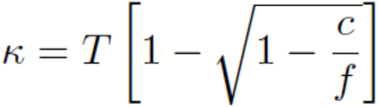

Therefore we take samples distributed as the square root of the iterations *t*.

### Software implementations

Mandrake is written in a combination of C++, CUDA, python and javascript. One of the major changes from the reference implementation of SCE is that we provide python bindings to the SCE method using pybind11 (29). The C++/CUDA part of mandrake which runs the entropic affinity preprocessing and modified SCE algorithm can be imported into any python program and called with ‘triplet’ sparse matrix data.

### Command line interface (python)

The full mandrake executable is available as a python executable which includes genetic distance calculation, and plotting of the output. We include numerous progress meters for each stage of computation, as on large datasets estimating time or eliminating computationally impossible steps is a necessity. The package can be installed using conda, and we provide online documentation and examples at https://mandrake.readthedocs.io/en/latest/.

### Optimisation of GPU code (CUDA)

We optimised the CUDA code through multiple rounds of profiling, the results of which can be accessed with the datasets on Zenodo. Briefly, this resulted in the following changes:

- Use of a callback function to output the objective function at each iteration, so convergence can be monitored.
- Use of CUDA graphs to run each iteration, which eliminates overheads from calls to the CUDA API at every step.
- Reversing the strides of the embedding Y from row-major to column-major, which can sometimes coalesce memory accesses. Changing the strides back (to be compatible with numpy) is done in a new device kernel.
- Use of parallel reductions from the cub library to calculate the objective at the end of each step.
- Use of the wrapper classes from the dust package to manage device memory (23).
- Elimination of thread divergences within warps.
- Inclusion of 32-bit and 64-bit versions of the code (64-bit operations are slower and use more registers, and some devices can only emulate 64-bit floating point operations, which can decrease performance greatly).
- Storing each worker’s random number generator state in registers, rather than writing to/from global memory whenever it is changed.
- Added compiler optimisations and loop unrolling.

### Static web app (WebAssembly and Javascript)

We optimised a version of Mandrake for the web (https://gtonkinhill.github.io/mandrake-web). This is particularly important to improve accessibility for users who have less experience running and installing bioinformatics programs on the command line. We made use of the Emscripten compiler to convert a slightly modified version of the C++ code used in the python package to WebAssembly, which executes within the browser on the user’s machine. This provides significant performance benefits over a pure javascript based implementation, and allows the web application to achieve similar speeds as the command line version on small to medium sized datasets. As the support for multi-threading in WebAssembly is still experimental, the web application currently only supports runs on a single CPU, so the command line version is still recommended for very large datasets.

The static web application was created using the Hugo site generator and custom javascript to interact with the compiled WebAssembly functions. A significant benefit of this approach is that once the website is loaded, there is no reliance on an internet connection and the entire analysis is run on the user’s local machine. This ensures that the user’s data is secure, as it is never uploaded, and which can be particularly useful in locations with poor internet connections where the uploading of any large dataset would be infeasible. It is also possible to run mandrake-web entirely offline.

### Data and code availability

Code: https://github.com/johnlees/mandrake and https://github.com/gtonkinhill/mandrake-web

Documentation: https://mandrake.readthedocs.io/en/latest/

Archived code: https://dx.doi.org/10.5281/zenodo.5579270

Commands used for analysis: https://github.com/gtonkinhill/mandrake_manuscript

Datasets: https://dx.doi.org/10.5281/zenodo.5572316

## Results

### Overview of Mandrake’s design

Figure 1 gives a graphical overview of the steps we use in mandrake to create a low dimensional embedding from genomic data. Pairwise genetic distances **X** between all samples are calculated from the genome data. Each element of **X**, of which there are *N*^2^, requires comparison of *M* genomic features. This is typically the largest calculation in mandrake, and we have highly optimised it and allow it to take advantage of many CPU cores where available. This makes calculation of distance matrices from up to millions of samples feasible. Each sample is reduced to the k-nearest neighbour distances on-the-fly to save space in memory. Note that although Figure 1 removes identical distances for visual clarity, in our code we retain them. We then use entropic affinity, as described in the introduction, to convert these distances into a conditional probability distribution, as described in the introduction. Figure 1 shows an example for sample *x*_3_, which has nearest neighbours *x*_1_ at one SNP away and *x*_2_ at two SNPs away. These are converted into probabilities using the height of Gaussian as shown, with a variance found to match the chosen perplexity through interval bisection.

**Figure 1:**
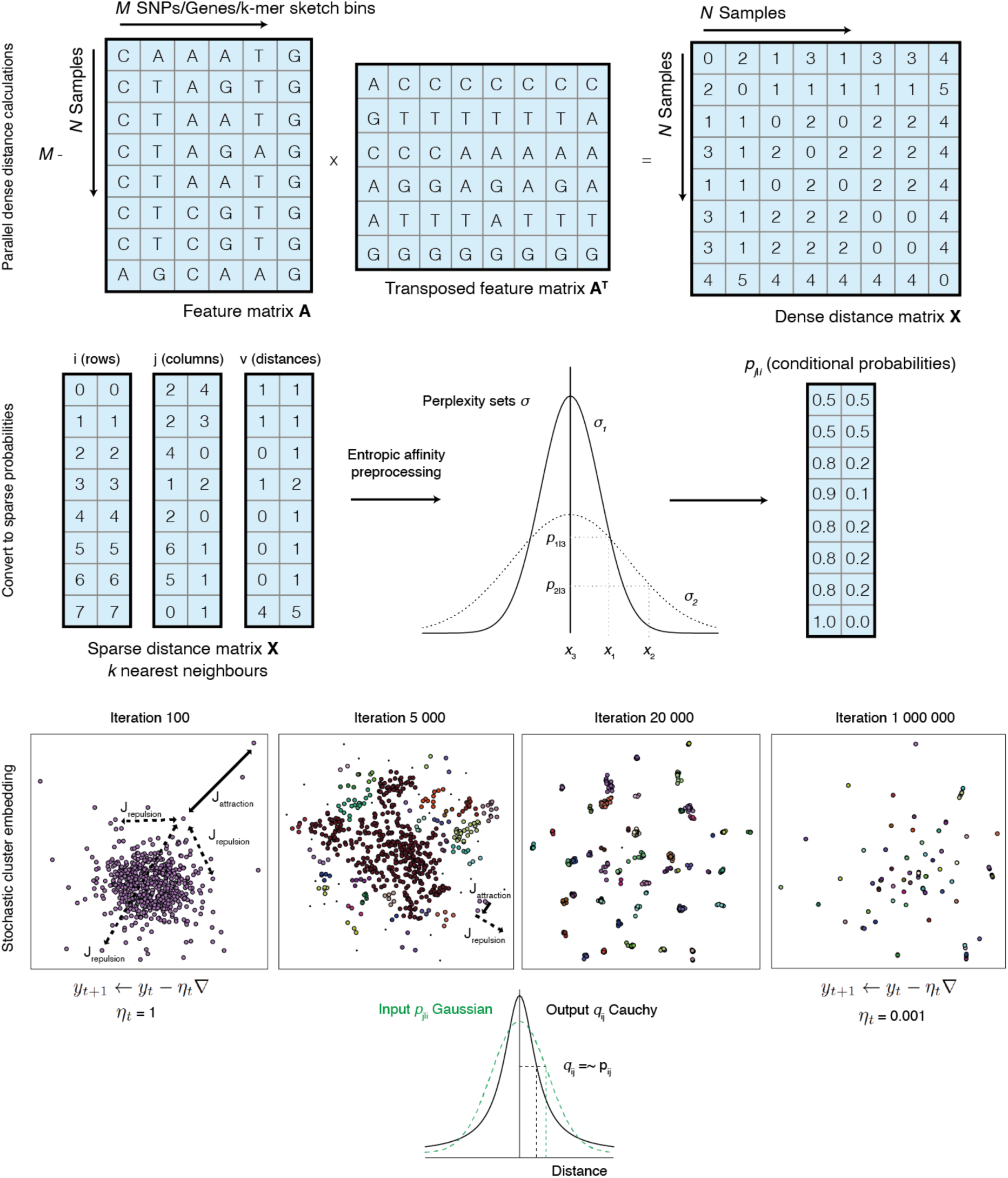
Overview of the mandrake software. Firstly, a genomic dataset, which may be a multiple sequence alignment, gene presence/absence or sequence sketches is used to calculate all pairwise distances between samples. Each entry in the distance matrix **X** is then the number of different features between each pair of samples. Each row of **X** is sorted, and the lowest *k* values (excluding self-matches of zero) are retained in triplet format. Entropic affinity converts these sparse distances to conditional probabilities, which can be thought of as the probability of selecting sample *x*_j_ as a neighbour, if probabilities are normally distributed. The user sets a perplexity parameter, which is used to set the variance of the distribution for each sample. Stochastic cluster embedding is run over a user-set number of iterations of stochastic gradient descent.

Stochastic cluster embedding is then run; we make the user specify the number of iterations to run for and do not stop until this is reached. Some example frames across the SGD iterations are shown. At the start, points in the lower dimensional space are randomly distributed, but are moved around more as the learning rate is higher. Later on, points are in clusters, and move smaller amounts along their gradient each step due to the lower learning rate. Some example attractive *J*_attraction_ and repulsive *J*_repulsion_ gradient steps are shown on the first two panels. Points are selected for attraction more frequently if they have a higher conditional probability. This has the effect that within a cluster (close in the higher dimension; higher conditional probabilities) points are pushed together. Repulsion is between any pair of points, which at later stages of the algorithm repulses clusters from one another, with the attractive force keeping the cluster together.

Applying SGD to *D* tries to make the input distribution (the conditional probabilities *p*_*j*|*i*_) as similar as possible to the output distribution (set by a Cauchy distribution *q*_*ij*_). This is shown at the bottom of figure 1 – an example with two points with the same input and output probability on the y-axis, are a small distance apart on the x-axis. Therefore, close distances in the higher dimensional space of **X** will also be close in the lower dimensional space **Y**. The heavy tails of the output Cauchy (black, solid distribution) apply less penalisation if smaller input probabilities are further apart than the tails of a Gaussian distribution (green, dashed distribution).

Our resulting implementation runs in a few hours, on up to around one million samples (table 1). Some variation in runtimes not directly proportional to the number of iterations is observed, this is typically due to setting a number of workers to aim for a maximum 10% clash rate on each dataset, such that efficiency increased in the larger datasets. Our web app is responsive up to the range of 10,000 samples, which completes in around a minute, depending on the input data type.

**Table 1:** Resource usage for mandrake on datasets used. The first row shows use of the static web app on a single core. All other reported times used 60 CPU cores, and where applicable an Nvidia 3090 RTX GPU was with double precision (fp64) or single precision (fp32). Note, GPU distance calculations are supported for sketches, but the table reports the CPU time only. The HIV pol gene alignment used was a random 5,000 subset from the Los Alamos public database (30).

| Dataset | Number of samples | Distance calculation | Maximum memory | Iterations used | SCE time (CPU) | SCE time (GPU) |
| --- | --- | --- | --- | --- | --- | --- |
| HIV-1 *pol* gene alignment | 5,000 | 35 s (web) | NA | $1 \times 10^6$ | 11s (web) | NA |
| *S. pneumoniae* accessory genome | 20,047 | 3 mins | 6.5 Gb (host) / 0.35 Gb (GPU) | $3 \times 10^8$ | 7.8 mins | 5s (fp64) |
| SRA bacterial assemblies | 661,406 | 3 hrs | 7 Gb (host) / 3 Gb (GPU) | $5 \times 10^{11}$ | 186 hrs | 1.3hrs (fp64) |
| SARS-CoV-2 alignment | 941,981 | 101 hrs | 26 Gb (host) / 13 Gb (GPU) | $1 \times 10^{12}$ | 372 hrs | 2.8 hrs (fp64) / 2.3 hrs (fp32) |

Before interpreting our results on different datasets, we recap some key features of non-linear embeddings (31):

- Cluster sizes in the embedding space do not relate to the number of points in the cluster, or its genetic diversity. In SCE particularly, many points will be heavily overplotted, and the density plot should be used for determining the number of samples in one region.
- Distances between clusters do not correspond to their genetic distances. Two well-separated clusters, close together, are not necessarily more genetically similar than two well-separated clusters at opposite ends of the plot.
- Perplexity can greatly affect results, and runs at a few different perplexities should typically be attempted. At lower perplexities, structure can sometimes be found where there is none. Complex topological relationships are generally not expected in genetic data, but where these may exist (in the presence of extensive horizontal gene transfer) multiple perplexity runs may be able to find these.

### Clustering the accessory genome of 20k

#### Streptococcus pneumoniae

To demonstrate the ability of the SCE embedding to identify meaningful clusters within a large dataset we first considered a collection of 20,047 *S. pneumoniae* genomes which consisted of a subset of high quality genomes from the Global Pneumococcal Sequencing project and two other pneumococcal genome surveillance studies (4,32–34). *S. pneumoniae* is a highly recombinant bacterial species with an extensive accessory genome that has been shown to be highly structured (35,36). This makes it a good example for investigating the ability of Mandrake to identify clusters from a gene presence/absence matrix.

We first inferred a pangenome gene presence/absence matrix using Panaroo v1.2 (37). This resulted in a binary matrix consisting of 27,322 features (genes) and 20,047 genomes which was used as input to Mandrake.

Figure 2 and supplementary video 1 indicate the resulting embedding with points coloured according to which of the Global Pneumococcal Sequencing Clusters (GPSCs) each genome belonged to (4,32). Those clusters with less than 50 genomes are coloured in grey with the Mandrake embedding placing them together in a single large group. This is similar to the behaviour of other clustering algorithms such as BAPS where outlying genomes are often grouped together into a single cluster representing the broader genomic background (30,38). To compare the observed clustering in the 2D embedding to the underlying GPSCs we calculated the rand index after first clustering the embedded points using HDBSCAN (28). The Rand index is a measure of similarity between the two clusterings and gives an indication of the accuracy with which one clustering predicts the other. The Mandrake embedding was found to have an index of 0.987 which was similar but still higher than that found using common alternative embeddings including UMAP, t-SNE and PCA (Supplementary Table 1). This suggests that Mandrake is able to produce a biologically meaningful embedding quickly using the presence and absence of accessory genes as input.

**Figure 2:**
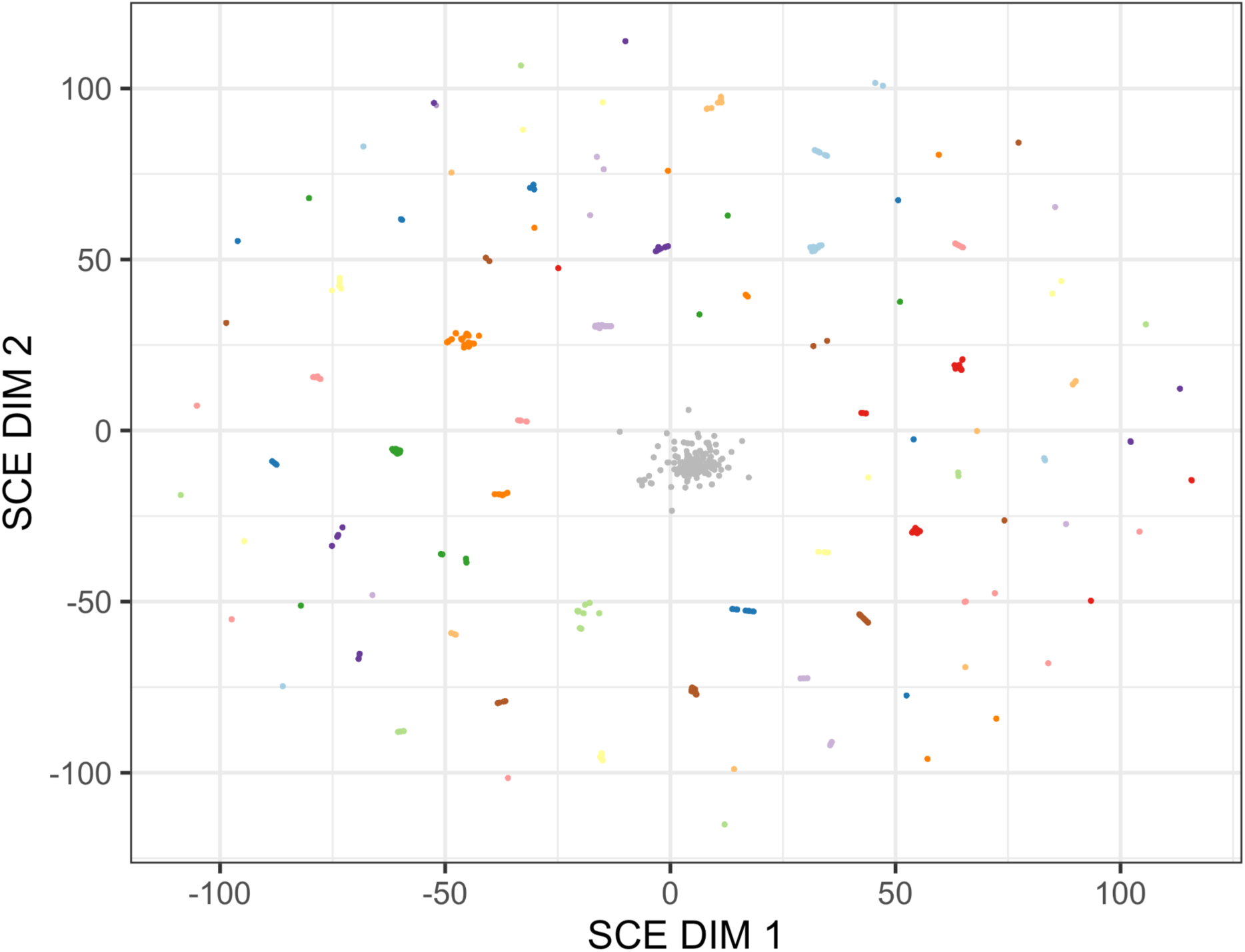
A 2-dimensional embedding calculated using Mandrake from the accessory gene presence/absence matrix of 20,047 pneumococcal genomes. Points are coloured by the underlying Global Pneumococcal Sequence Cluster to which they belong. Those GPSCs with less than 50 isolates are coloured in grey.

#### Clustering ∼650k bacterial genome assemblies from public databases

We then used SCE to search for structure across the space of highly diverse bacterial genomes. A recent analysis produced curated and assembled bacterial samples from the SRA database, producing 661,406 high-quality bacterial genomes (39). We downloaded these assemblies from ftp.ebi.ac.uk/pub/databases/ENA2018-bacteria-661k (39) and sketched their 14-mers. We then used these sketches to calculate Jaccard distances between 14-mers, which we used to produce a sparse matrix with the 100 nearest neighbours. Using this, we ran mandrake for 5×10^11^ iterations using 65,536 workers on a GPU, which took two hours. The objective stabilised around halfway through the run, and the resulting embedding can be seen in Figure 3, and an animation of the SCE iterations in supplementary video 2.

**Figure 3:**
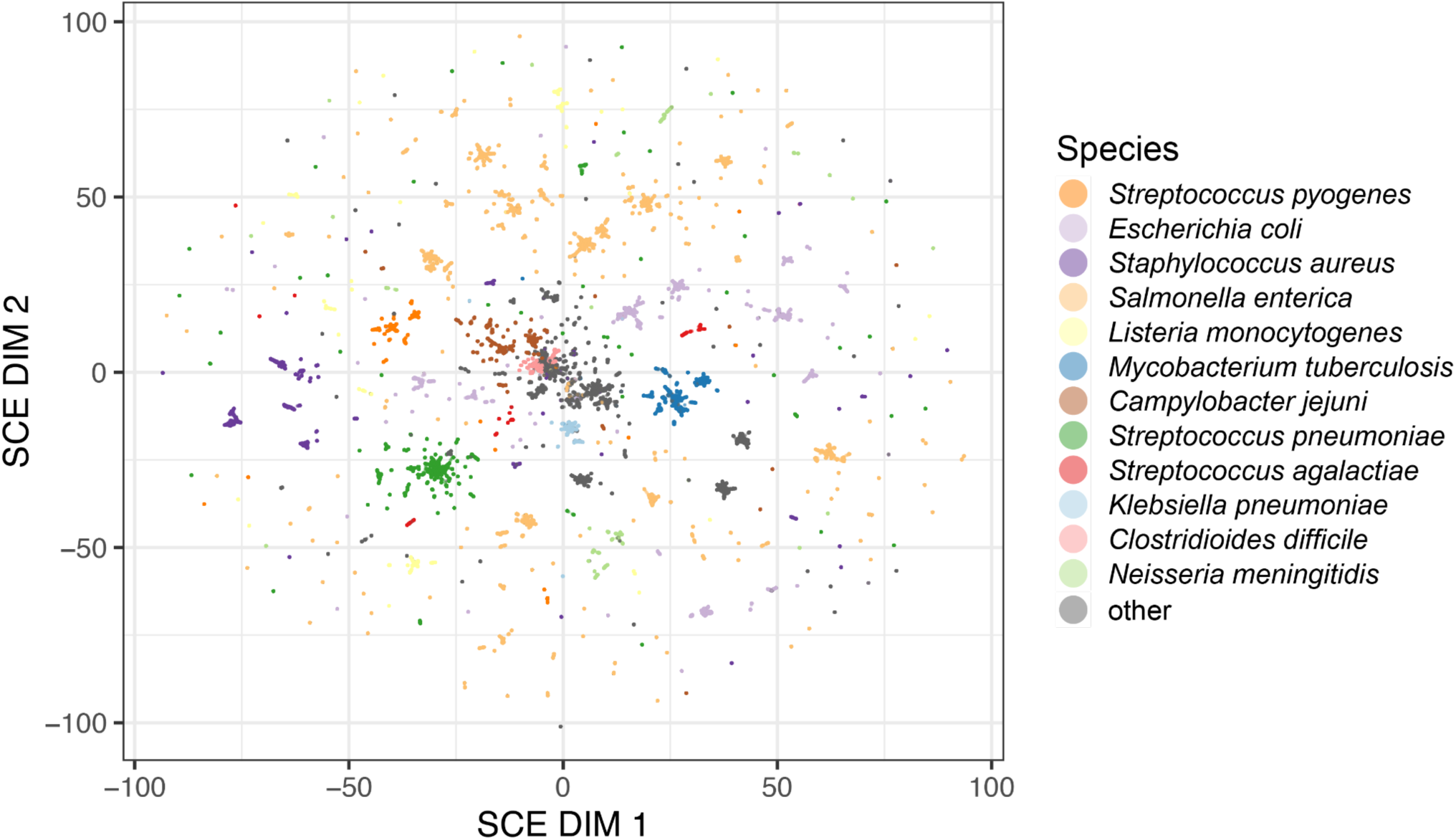
A Mandrake embedding of 662,406 bacterial genomes from the SRA with Jaccard distances calculated using sketches of 14-mers. Species with at least 10,000 genomes in the data set are coloured.

We found that the most common species formed clear clusters in the embedding space, with the exception of *Closteroides difficile* in the centre of the space, which overlaps with many other gut pathogens. This is likely due to be from gut samples where the sequence contained multiple species, but just the most abundant species was reported. Most species were split into multiple clusters, likely representing strains within species, or subspecies (in the case of e.g. *Salmonella enterica* which appears in multiple clusters over the whole embedding space). Some other interesting examples include *Listeria monocytogenes* which has genetically distinct major lineages (40), and appears as separate clusters spread around the embedding. *Mycobacterium tuberculosis* is split into 5-10 clusters, which are close together in the embedding, and have a larger radius than e.g. *Escherichia coli* clusters, despite harbouring much less genetic diversity. The non-linear nature of the embedding is therefore also able to capture structure across a range of genetic scales. This also demonstrates the points of cluster size not having a direct interpretation, but between cluster distances sometimes retaining meaning.

#### Clustering ∼1M SARS-CoV-2 genome alignments from public databases

We next considered Mandrake’s ability to embed highly similar genomes into clusters by running the algorithm in its multiple sequence alignment mode on a cleaned subset of 941,981 SARS-CoV-2 genomes downloaded from the ENA (covid19dataportal.org). Of the original 977,048 genomes downloaded, we filtered out 35,067 which had a length less than 90% of the Wuhan.1 reference genome or were made up of more than 5% ambiguous nucleotide calls. Each genome was assigned to a SARS-CoV-2 lineage using Pangolin (5,41). After generating a multiple sequence alignment of the genomes using MAFFT v7.487 we ran Mandrake in it’s ‘alignment’ mode which calculates the pairwise hamming distance between genomes ignoring ambiguous base calls. Mandrake was run for 1×10^12^ iterations on a GPU with 94,976 workers, which took 3.7 hours. The resulting embedding is shown in Figure 4 and supplementary video 3, with the major SARS-CoV-2 lineages comprising more than 10,000 genomes assigned different colours. Interestingly, the major variants of concern including the Delta and Alpha lineages are clearly visible in the embedding indicating that Mandrake is able to identify biologically meaningful structure within very large but highly similar genomes.

**Figure 4:**
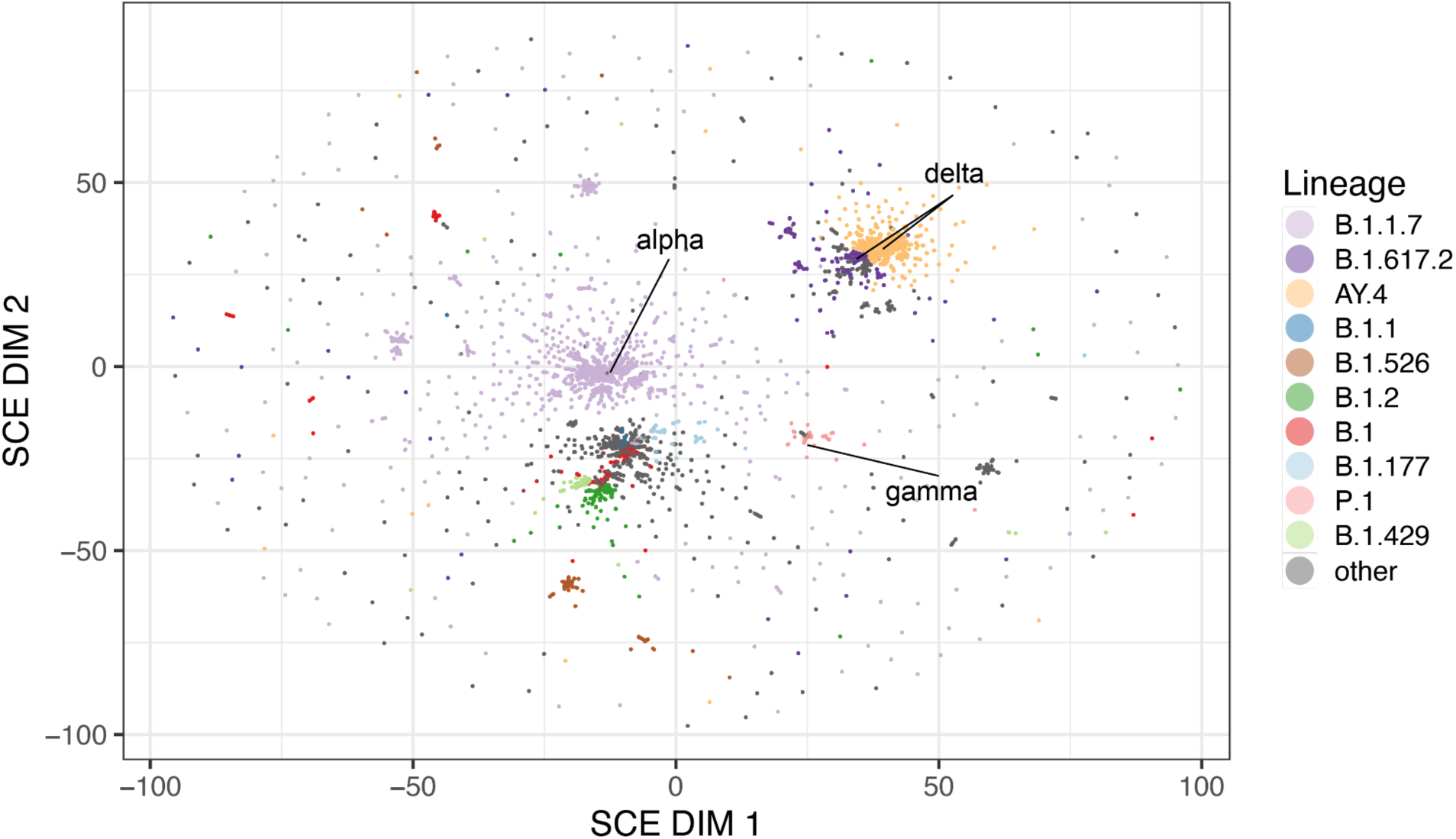
A Mandrake embedding of 941,981 SARS-CoV-2 genomes downloaded from the ENA. Points corresponding to genomes within large Pangolin lineages (>10,000 members) are coloured and variants of concern are manually labelled.

##### Discussion

Currently population genomics of pathogens is experiencing an unprecedented pace of genome sequencing on a global scale, which poses a challenge to many standard workflows for data analysis. As many downstream epidemiological or evolutionary analyses work within identified clusters, the first tasks in population genomics workflows are to understand the population structure and the extent of clustering of the input genomes, both of which are difficult to do with increasing data size. While some new highly scalable methods have been developed for this task, they are frequently species-specific (42). Doing this in a manner which can be visualised is particularly helpful, especially given the high dimensions and complex relationships inherent in genomic datasets.

In this article we have presented the mandrake software, which meets these particular needs and offers programmatic plotting options and interactive exploration of the data. Our current software architecture scales well to even the largest available contemporary pathogen genomics data sets. However, in future it would clearly benefit from reducing the quadratic computational complexity of the input genome distance calculations, which could for example be achieved with subsampling of the data by picking representative samples among highly similar genomes. This has been achieved in other packages by assuming genetic distances generally obey the triangle inequality (18,43).

Another interesting opportunity for further research and development stems from the challenge of optimizing output plots using models of human perception. Here, we used user-guided training in SCE to determine a parameter *s*, which governs the display of clusters in the output embedding. Recent results in perception modeling for visualization have demonstrated notable improvements over default software options for scatterplots, where optimized designs can much better adapt to an increase in data density (44). There are multiple display parameters that could be adjusted in order to give a human expert an enhanced view into the data structure, such as the marker size, their opacity and colors. Optimization of such parameters using models of human cognition has the potential to resolve the visualization challenges arising from extremely high dimensionality of the data, not only for cluster embedding as considered here, but also for other complex objects such as phylogenetic trees. An exciting opportunity for considering this untapped potential arises from combining cognitive models with recent developments in Approximate Bayesian Computation (ABC), which allows efficient fitting of simulator-based models to data when the likelihood calculations are intractable (45,46). Several successful examples of using ABC in cognitive model inference (47,48) suggest that this approach could be fruitful for making improved displays of high-dimensional population genomic data.

Nevertheless, we have been able to make mandrake useful across a range of scales and to a range of users, scaling from within-browser analysis, through multicore CPU use on the command line, up to high power graphics cards. The functions provided may also serve as a basis for the analysis of pairwise relationships between genomic data in other tools, such as phylogenetics (49), selection analysis (50) and mathematical modelling (51).

## Acknowledgements

We would like to thank the many researchers who generated genomic data and then openly deposited it in databases free of usage restrictions (SRA and INSDC), without whom developing methods to analyse large microbial datasets would be considerably more challenging. In particular, we are grateful to the GPS consortium, Grace Blackwell and the COVID-19 data portal for making the genomic data and associated metadata from their analyses freely available and easy to access in a reproducible and reusable manner. JAL acknowledges funding from the MRC Centre for Global Infectious Disease Analysis (reference MR/R015600/1), jointly funded by the UK Medical Research Council (MRC) and the UK Foreign, Commonwealth & Development Office (FCDO), under the MRC/FCDO Concordat agreement and is also part of the EDCTP2 programme supported by the European Union. GTH was funded by the Norwegian Research Council grant no. 299941. JC was funded by the European Research Council grant no. 742158.

## Figure and table captions

**Supplementary Table 1:** Rand index between PopPUNK clusters (GPSCs) on the S. pneumoniae accessory data, and HDBSCAN clusters made from four different embedding methods.

| Method | Rand Index |
| --- | --- |
| Mandrake | 0.98697 |
| UMAP | 0.98660 |
| tSNE | 0.98598 |
| PCA | 0.84303 |

**Supplementary video 1:** An animation of mandrake running on the *S. pneumoniae* accessory genome data, with the same parameters as figure 1.

**Supplementary video 2:** An animation of mandrake running on the SRA bacterial genomes data, with the same parameters as figure 2.

**Supplementary video 3:** An animation of mandrake running on the SARS-CoV-2 genomes data, with the same parameters as figure 3.

